# Selene: a PyTorch-based deep learning library for biological sequence-level data

**DOI:** 10.1101/438291

**Authors:** Kathleen M. Chen, Evan M. Cofer, Jian Zhou, Olga G. Troyanskaya

**Author notes:** These authors contributed equally to this work (Kathleen M. Chen, Evan M. Cofer). To whom correspondence should be addressed. Olga G. Troyanskaya.

## Abstract

To enable the application of deep learning in biology, we present Selene (https://selene.flatironinstitute.org/), a PyTorch-based deep learning library for fast and easy development, training, and application of deep learning model architectures for any biological sequences. We demonstrate how Selene allows researchers to easily train a published architecture on new data, develop and evaluate a new architecture, and use a trained model to answer biological questions of interest.

## Main

Deep learning describes a set of machine learning techniques that use stacked neural networks to extract complicated patterns from high-dimensional data^1^. These techniques are widely used for image classification and natural language processing and have led to very promising advances in the biomedical domain, including genomics and chemical synthesis^1–3^. In regulatory genomics, networks trained on high-throughput sequencing data (e.g. ChIP-seq), or “sequence-based models,” have become the *de facto* standard for predicting regulatory and disease impact of mutations^4–7^. While deep-learning related publications are often accompanied by the associated pre-trained model^6,8,9^, a key challenge in both developing new deep learning architectures and training existing architectures on new data is the lack of a comprehensive, generalizable, and user-friendly deep learning library for biology.

Beyond regulatory genomics, sequence-level deep learning models have broad promise in a wide range of research areas, including recent advances on prediction of disease risk of missense mutations in proteins^10^ and potential applications to, for example, predicting target site accessibility in genome editing. We must enable the adoption and active development of deep learning-based methods in biomedical sciences. For example, a biomedical scientist excited by a publication of a model capable of predicting the disease-associated effect of mutations should be able to train a similar model on their own ChIP-seq data focused on their disease of interest. A bioinformatician interested in developing new model architectures should be able to experiment with different architectures and evaluate all of them on the same data. Currently, this requires advanced knowledge specific to deep learning^2,11^, substantial new code development, and associated time investment far beyond what most biomedical scientists are able to commit.

Here we present Selene, a framework for developing **se**quence-level deep **le**arning **ne**tworks, that provides biomedical scientists with comprehensive support for model training, evaluation, and application across a broad range of biological questions. Sequence-level data refers to any type of biological sequence such as DNA, RNA, or protein sequences and their measured properties (e.g. TF binding, DNase sensitivity, RBP binding). Selene contains modules for (1) data sampling and training for model development (Fig. 1a), and (2) prediction and visualization for analyses using the trained model (Fig. 1b, c). With Selene, researchers can run model development and analysis workflows out-of-the-box. For more advanced use cases, Selene provides templates for extending modules within each workflow so that users can adapt the library to their particular research questions.

**Figure 1.**
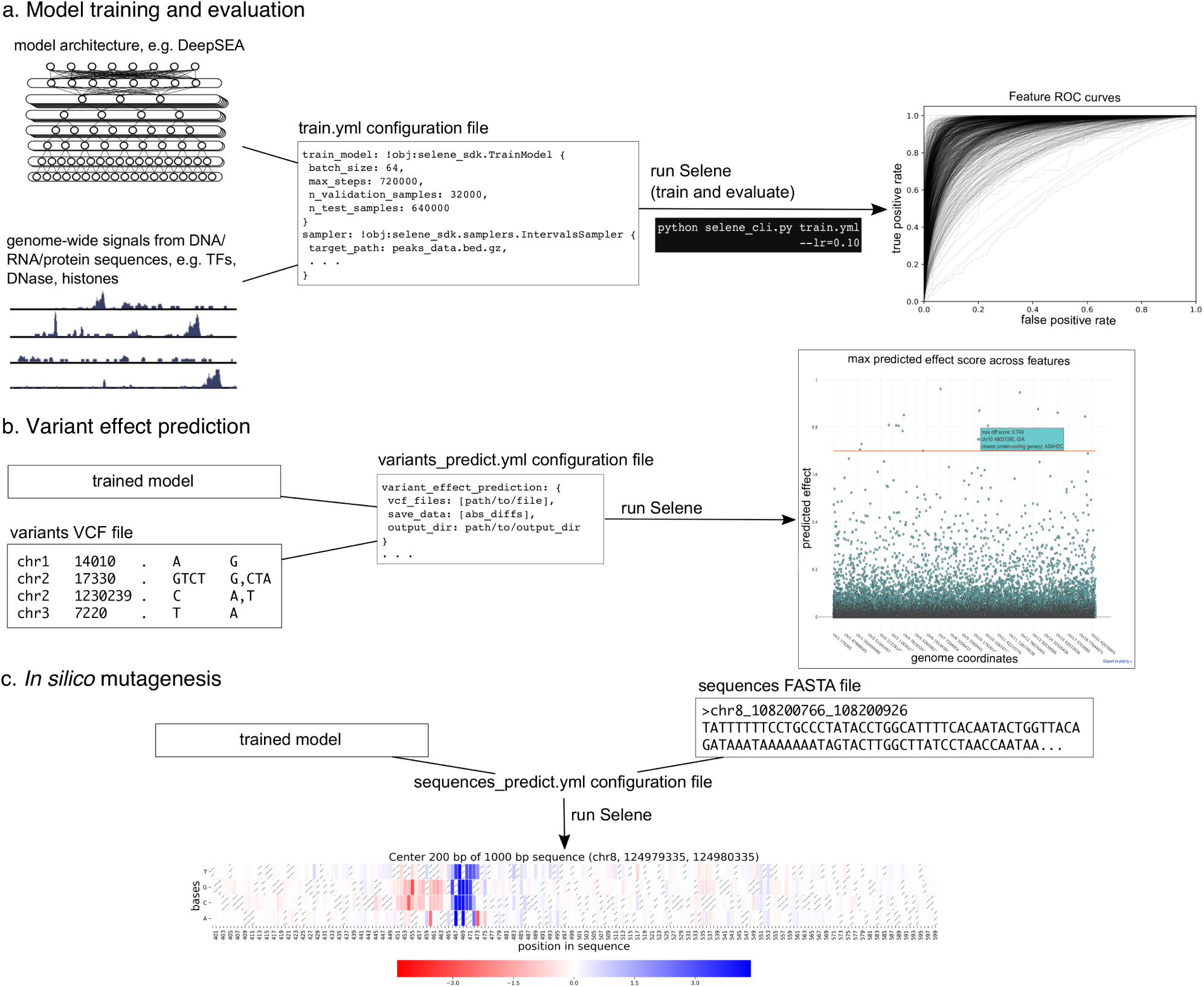
Overview of functionality provided in Selene. **(a)** Selene enables users to train and evaluate new deep learning models with very few lines of code. As input, the library accepts (left) the model architecture, dataset, and (mid) a configuration file that specifies the necessary input data paths and training parameters; Selene automatically splits the data into training and validation/testing, trains the model, evaluates it, and (right) generates figures from the results. **(b)** Selene also supports the use of trained models to interpret variants. In addition to being able to run variant effect prediction with the same configuration file format, Selene includes a visualization of the variants and their difference scores as a Manhattan plot, where a user can hover over each point to see variant information. **(c)** Users interested in finding informative bases that contribute to the binding of a certain protein can run Selene to get mutation effect scores and visualize the scores as a heatmap.

There has been recent work to make deep learning in biology more accessible: DragoNN is a toolkit for teaching deep learning in regulatory genomics; pysster^12^ is a Python package for training convolutional neural networks on biological sequence data; and Kipoi^13^ is a framework to archive, use, and build on published predictive models in genomics. These resources constitute the nascent software ecosystem for sequence-level deep learning. Selene is our contribution to this ecosystem. Selene supports general model development not constrained to a particular architecture (in contrast to pysster) or task (in contrast to DragoNN) and is designed for users with different levels of computational experience. Users are supported in tasks ranging from simply applying an existing model, to retraining it on new data (tasks also supported by Kipoi), to developing new model architectures (a task that is challenging to do with any other tool). The models developed using Selene can be shared and used through the Kipoi framework.

To demonstrate Selene’s capabilities for developing and evaluating sequence-level deep learning models, we use it to (1) train a published architecture on new data, (2) develop, train, and evaluate a new model (improving a published model) and (3) apply a trained model to data and visualize the resulting predictions.

### Case 1: Training an existing (e.g. published) architecture on a different dataset

In this case study, a researcher reads a manuscript about a deep learning model and wants to train the model on different data. For illustration, we will use the DeepSEA model architecture as a starting point in our case studies; however, Selene is completely general and a user can easily use or specify any model of their choice using modules in PyTorch.

Suppose a cancer researcher is interested in modeling the regulatory elements of the transcription factor GATA1, specifically focusing on proerythroblasts in bone marrow. This is a tissue-specific genomic feature that DeepSEA does not predict. The researcher downloads peaks data from Cistrome^14^ and a reference genome FASTA file. Once a researcher formats the data to match the documented inputs (https://selene.flatironinstitute.org/overview/cli.html) and fills out the necessary training parameters (e.g. batch size, learning rate), they can use Selene to train the DeepSEA architecture on their data with no new lines of Python code. In this example, they find that the model obtains an AUC of 0.942 on this feature (Fig. 2a).

**Figure 2.**
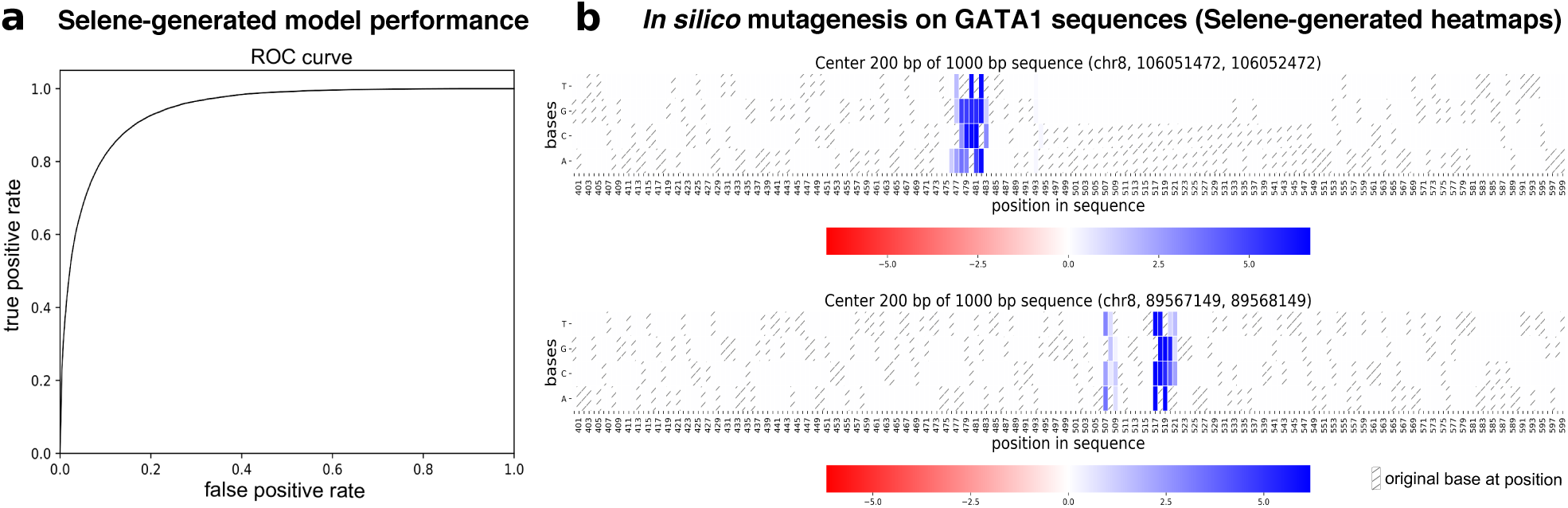
Figures generated by Selene. **(a)** Selene visualization of the performance of the model trained in case study 1. **(b)** Selene visualization of *in silico* mutagenesis on the case-study-trained model for 20 randomly selected GATA1 sequences in the test set (2 representative plots displayed here, all heatmaps generated are displayed in the example Jupyter notebook: https://github.com/FunctionLab/selene/blob/master/manuscript/case1/3_visualize_ism_outputs.ipynb). Bases in the original sequence are distinguished by the gray stripes in the heatmap cells.

Selene automatically generates training, testing, and validation samples from the provided input data. The samples generated for each partition can be saved and used in subsequent model development so that comparisons can be made across models with different architectures and/or parameters. Further, Selene automatically evaluates the model on the test set after training and, in this case, generates figures to visualize the model’s performance as receiver operating characteristic and average precision curves.

Now that the researcher has a trained model, they can use Selene to apply *in silico* mutagenesis to a set of GATA1 sequences from the test set. *In silico* mutagenesis involves “mutating” every position in the sequence to every other possible base^4^ (DNA and RNA) or amino acid (protein sequences) and examining the consequences of these “mutations.” Selene supports visualizing the outputs of *in silico* mutagenesis as a heatmap and/or motif plot. By visualizing the log2 fold change for these sequences in a heatmap, the researcher can see that the model detects disruptions in binding at the GATA motif (Fig. 2b).

We provide the code and results for this example in Selene’s GitHub repository (https://github.com/FunctionLab/selene/tree/master/manuscript/case1). The case study demonstrates that Selene enables researchers to quickly get started with sequence-based deep learning; researchers can train and apply a model to their area of interest with minimal code and deep learning knowledge. Without Selene, a researcher would need to dedicate substantial time to preprocess their datasets and write code for training and evaluating their model.

#### Case 2: Developing a new architecture and comparing performance across architectures

In another use case, a researcher may want to develop and train a new model architecture. For example, a bioinformatician might want to modify a published model architecture to see how that affects performance. First, the researcher uses modules in PyTorch to specify the model architecture they are interested in evaluating; in this case study, they try to enhance the DeepSEA architecture with batch normalization and three additional convolutional layers. The researcher specifies parameters for training and the paths to the model architecture and data in a configuration file and passes this as input to the library’s command-line interface. Training is automatically completed by Selene; afterwards, the researcher can easily use Selene to compare the performance of their new model to the original DeepSEA model on the same chromosomal holdout dataset.

In this case study, the researcher finds that the deeper architecture achieves an average AUC of 0.938 (Fig. 3a) and an average AUPRC of 0.362, which is an improvement over the average AUC of 0.933 and AUPRC of 0.342 of the original 3-convolutional-layer model. The researcher can share this model with a collaborator (e.g. a human geneticist, see case study 3 below) and upload it to the Kipoi^13^ model zoo, a repository of trained models for regulatory genomics, with which Selene-trained models are fully compatible (see an example at https://github.com/FunctionLab/selene/tree/master/manuscript/case2/3_kipoi_export).

**Figure 3.**
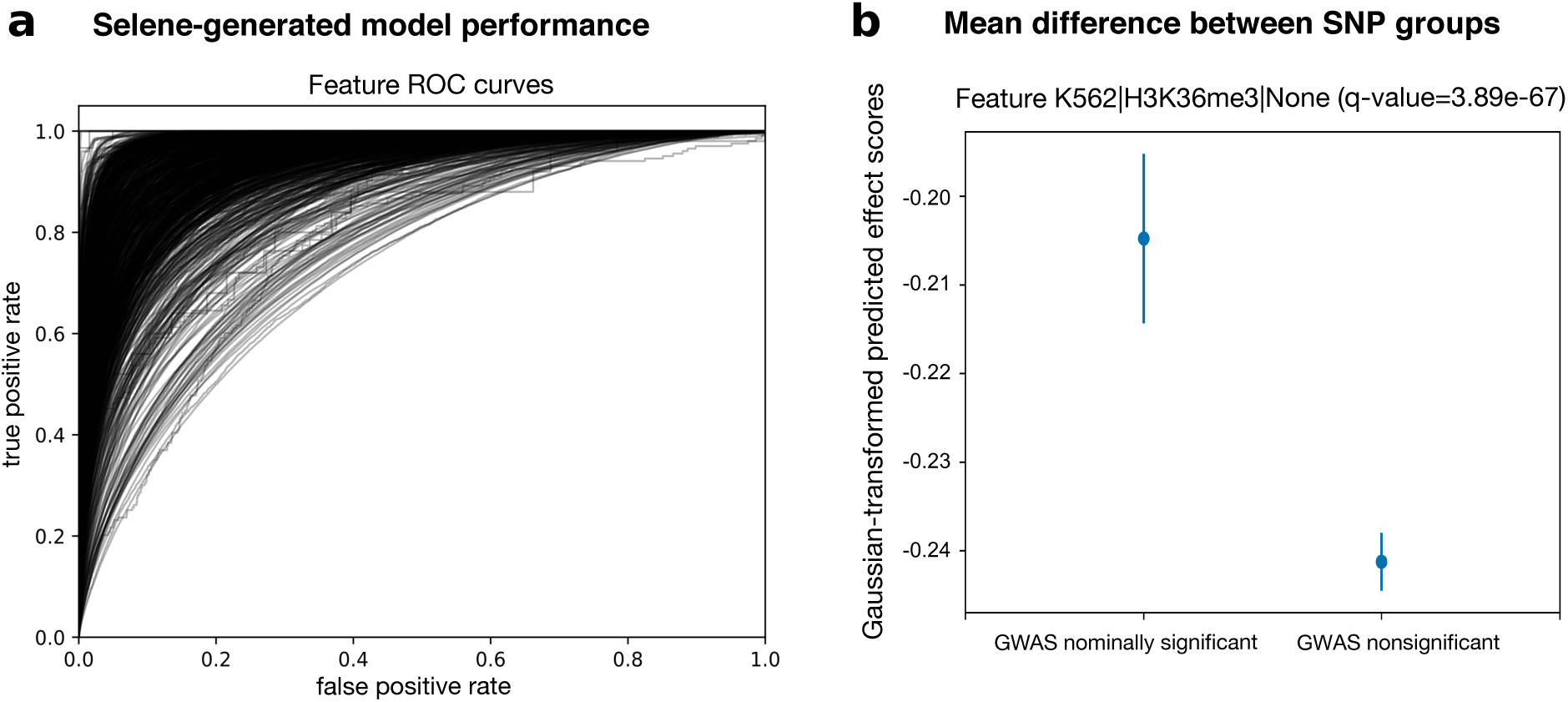
**(a)** Selene visualization of the performance of the trained model from case-study 2. **(b)** The trained model, on average, predicts greater differences for nominally significant variants (p-value < 0.05) in the IGAP early onset Alzheimer’s GWAS study compared to those that are nonsignificant (p-value > 0.50). Here we visualize the mean and 95% confidence intervals of the quantile-normalized (against the Gaussian distribution) predicted effect scores of the 2 variant groups for the genomic feature H3K36me3 in K562 cells, the feature in the model with the most significant difference (one-sided Wilcoxon rank-sum test, adjusted p-value using Benjamini—Hochberg of 3.89 × 10^-67^). After applying the multiple testing correction, 914 of the 919 features the model predicts showed a significant difference (a < 0.05) between the groups.

Using Selene, researchers can substantially reduce the amount of work needed to develop, train, and compare new models. Researchers are able to focus on experimenting with various model architectures rather than writing all new code for model training and evaluation.

#### Case 3: Applying a new model to variants

In this case study, a human geneticist studying Alzheimer’s wants to apply the model developed in case study 2 above, so they first assess its ability to prioritize potential disease-associated variants. Specifically, they use Selene to make variant effect predictions for nominally significant variants (p-value < 0.05) and nonsignificant variants (p-value > 0.50) in the International Genomics of Alzheimer’s Project^15^ Alzheimer’s disease GWAS^16^. The researcher finds that the predicted effect is significantly higher for GWAS nominally significant variants versus non-significant variants, indicating that the new model is indeed able to prioritize potential disease-associated variants (one-sided Wilcoxon rank-sum test, most significant feature H3K36me3 in K562 cells has an adjusted p-value, by Benjamini—Hochberg correction, of 3.89 × 10^-67^) (Fig. 3b).

Selene’s modeling capability extends far beyond case studies shown above. The library can be applied to not only DNA, but also RNA and protein sequences; and not only chromatin data, but any current genome-, transcriptome-, or even proteome-wide measurements. We developed Selene to increase the accessibility of deep learning in biology and facilitate the creation of reproducible workflows and results. Furthermore, Selene is open-source software that will continue to be updated and expanded based on community and user feedback.

## Methods

### Overview of Selene

Selene consists of two components: a Python library for developing sequence-level neural networks, and a command line interface for prototypical use cases of the library (i.e. training a new model, evaluating an existing model, and analyzing sequence data and variants with a trained model). We herein refer to these components as the software development kit (SDK) and the command line interface (CLI) respectively. Importantly, all functionality provided by the CLI is also available to the user through the SDK. Rather than supplanting the SDK, the CLI is intended to maximize code reuse and minimize user time spent learning SDK by heavily reducing the configuration tasks left to the user (e.g. when GPU usage is specified, the CLI ensures all appropriate computations are performed on the GPU). When appropriate, the SDK does deliver functionality beyond that of the CLI. For instance, the SDK includes several data visualization methods that would be too unwieldy as executables run from the command line.

Thorough documentation for the SDK is available at https://selene.flatironinstitute.org, and tutorials for both the CLI and SDK can be found at https://github.com/FunctionLab/selene/tree/master/tutorials. Notably, one tutorial demonstrates how to use Selene to train a deep neural network regression model (https://github.com/FunctionLab/selene/blob/master/tutorials/regression_mpra_example/regression_mpra_example.ipynb). This tutorial illustrates Selene’s use outside of the models of transcriptional regulation shown in the case studies.

### Selene software development kit

The Selene SDK, formally known as *selene_sdk*, is an extensible Python package intended to ease development of new programs that leverage sequence-level models through code reuse. The Selene CLI is built entirely upon the functionality provided by the SDK, but it is likely that users will use the SDK outside this context. For example, after training a new sequence-level model with the CLI, one could use the SDK in conjunction with a Python-based web application framework (e.g. Flask, Django) to build a web server so that other researchers can submit sequences or variants and get the trained model’s predictions as output.

Leveraging the SDK in a user Python project is no different from using any other Python module. That is, one only needs to import the *selene_sdk* module or any of its members, and supply them with the correct parameters. The runtime behavior of each component of *selene_sdk*, as well as the required parameters for all members of *selene_sdk*, is described in detail in the online documentation (https://selene.flatironinstitute.org/overview/overview.html).

### Selene command line interface

The Selene CLI is a usable program to be run from the command line by the user. It encapsulates the configuration, execution, and logging of Selene’s most common use cases. Said use cases are embodied by the CLI’s three commands: *train, evaluate*, and *analyze*. These commands are used to train new models, evaluate the performance of trained models, and analyze model predictions (perform *in silico* mutagenesis or variant effect prediction) respectively. Each command configures its specific runtime environment with a combination of command line arguments and parameters drawn from user-provided configuration files. The flexibility of these configuration files allows them to leverage user-developed code as well, and further extends the usability of the CLI. We provide a step-by-step tutorial that describes the CLI configuration file format and shows some example configuration keys and values at https://github.com/FunctionLab/selene/blob/master/tutorials/getting_started_with_selene/getting_started_with_selene.ipynb. Additional examples of CLI configuration files are available at https://github.com/FunctionLab/selene/tree/master/config_examples as well. Finally, comprehensive documentation detailing all possible configurations supported by Selene can be found at https://selene.flatironinstitute.org/overview/cli.html. Users can reference any of these resources when creating their own configuration files.

### Model Architectures

DeepSEA architecture used in case 1 (from the supplementary note in the DeepSEA publication^4^):

1. Convolutional layer (320 kernels, window size = 8, step size = 1)
2. Pooling layer (window size = 4, step size = 4)
3. Convolutional layer (480 kernels, window size = 8, step size = 1)
4. Pooling layer (window size = 4, step size = 4)
5. Convolutional layer (960 kernels, window size = 8, step size = 1)
6. Fully connected layer (919 genomic features)
7. Sigmoid output layer

Dropout proportion (proportion of outputs randomly set to 0):

Layer 2: 20%
Layer 4: 20%
Layer 5: 50%
All other layers: 0%

Architecture used in cases 2 and 3:

1. Convolutional layer (320 kernels, window size = 8, step size = 1)
2. Convolutional layer (320 kernels, window size = 8, step size = 1)
3. Pooling layer (window size = 4, step size = 4)
4. Convolutional layer (480 kernels, window size = 8, step size = 1)
5. Convolutional layer (480 kernels, window size = 8, step size = 1)
6. Pooling layer (window size = 4, step size = 4)
7. Convolutional layer (960 kernels, window size = 8, step size = 1)
8. Convolutional layer (960 kernels, window size = 8, step size = 1)
9. Fully connected layer (919 genomic features)
10. Sigmoid output layer

Dropout proportion:

Layer 5: 20%
Layer 8: 50%

Batch normalization applied after layers 2, 5, and 8 and before dropout.

Both architectures use the binary cross-entropy loss function and stochastic gradient descent optimizer (momentum = 0.9, weight decay = 10^-6^).

### Reproducing the case studies

Below, we have described the steps taken for each of the case studies. The code required to reproduce each case study is included in the GitHub repository (https://github.com/FunctionLab/selene/tree/master/manuscript). We have also created Zenodo records for each case that contain all the input data, data processing scripts, and outputs files generated from Selene:

- Case 1. doi:10.5281/zenodo. 1442433
- Case 2. doi: 10.5281/zenodo.1442437
- Case 3. doi: 10.5281/zenodo.1445555

#### Case 1: Training a state-of-the-art architecture on a different dataset

##### Steps to train DeepSEA on new data

1. Download the data from Cistrome. In this case, we are only working with 1 dataset for 1 specific genomic feature. Cistrome ID: 33545, measurements from GSM970258 (Xu et al., 2012): http://dc2.cistrome.org/api/downloads/eyJpZCI6IjMzNTQ1In0%3A1fujCu%3ArNvWLCNoET6o9SdkL8fEv13uRu4b
2. Format the data. We use tools from Samtools^17^ (specifically, tabix^18^ and bgzip from HTSlib, https://www.htslib.org/). Create a .bed file of chromosome, start, end, and the genomic feature name (useful when there is more than 1 feature). Sort this file and compress it into a .gz file. Tabix-index this file. Specific commands:

a. Only use the columns [chr, start, end]:

~~~
cut -f 1–3 **<peaks-file>** > **<peak-coordinates-file>**
~~~ Note: Eventually, we will add support for parsing BED files with strand specific features and/or continuous values that quantify these features.
b. Add the genomic feature name as the 4th column of the file:

~~~
sed -i “s/$/\t**<feature-name>**/” **<peak-coordinates-file>**
~~~
c. Sort the file by [chr, start, end]:

~~~
sort -k1V -k2n -k3n **<peak-coordinates-file>** > **<sorted-coordinates-file>**
~~~
d. Compress the file:

~~~
bgzip **<sorted-coordinates-file>**
~~~ This compresses the file to a .gz file in-place. To separately generate the .gz file, run

~~~
bgzip -c **<sorted-coordinates-file> > <sorted-coordinates-file>**.gz
~~~
e. Tabix index the file:

~~~
tabix -p bed **<sorted-coordinates-file>**.gz
~~~
3. Create a file of distinct features that the model will predict, where each feature is a single line in the file. This can easily be created from the .bed file in step 2 by running:

~~~
cut -f 4 **<peak-coordinates-file>** | sort -u > **<distinct-features>**
~~~
4. Download the GRCh38/hg38 FASTA file. We downloaded the reference sequences GRCh37/hg19 and GRCh38/hg38 used in our analyses from ENCODE: https://www.encodeproject.org/data-standards/reference-sequences/.
5. Specify the model architecture, loss, and optimizer as a Python file. **This is done for you in the case of DeepSEA**: https://github.com/FunctionLab/selene/blob/master/models/deepsea.py
6. Fill out the configuration file with the appropriate file paths and training parameters. We recommend starting from one of the example training files in the tutorials (e.g. https://github.com/FunctionLab/selene/blob/master/tutorials/getting_started_with_selene/getting_started_with_selene.ipynb) or in the “config_examples” directory (https://github.com/FunctionLab/selene/tree/master/config_examples). You can also review the documentation for the configuration parameters on Selene’s website (https://selene.flatironinstitute.org/overview/cli.html).
7. Run Selene.

##### Steps to apply and visualize the results of *in silico* mutagenesis

1. Collect sequences you want to visualize as a FASTA file. **For this particular case, we provide a script to do so** (https://zenodo.org/record/2214130/files/data.tar.gz).
2. Fill out the configuration file with the appropriate file paths (path to the FASTA file, information about the trained model).
3. Run Selene. You will get the raw predictions and the log2 fold change scores as output files.
4. Follow one of the Jupyter notebook tutorials for *in silico* mutagenesis (https://github.com/FunctionLab/selene/tree/master/tutorials) to generate visualizations for the sequences. **We have done this in** https://github.com/FunctionLab/selene/blob/master/manuscript/case1/3_visualize_ism_outputs.ipynb.

#### Case 2: Developing a new architecture and making model comparisons

##### Steps to train “deeper DeepSEA” on the same exact data as DeepSEA

1. Download the code and data bundle from the DeepSEA website (http://deepsea.princeton.edu/media/code/deepsea_train_bundle.v0.9.tar.gz). You only need the .mat files in this directory. We also include a file listing the 919 genomic features that the model predicts. This is from the resources directory in the standalone version of DeepSEA (http://deepsea.princeton.edu/media/code/deepsea.v0.94b.tar.gz). **Zenodo record:** https://zenodo.org/record/2214970/files/DeepSEA_data.tar.gz.
2. Fill out the configuration file for Selene’s MultiFileSampler (https://selene.flatironinstitute.org/overview/cli.html#multiple-file-sampler) and specify the path to each .mat file for training, validation, and testing.
3. Run Selene.

Please see the DeepSEA publication for details about data processing and training: https://www.nature.com/articles/nmeth.3547#methods.

In the main text, we report test performance for the model trained using the online sampler.

When training on the same exact data (the .mat files) as DeepSEA, we achieve an average AUC of 0.934 and an average AUPRC of 0.361.

##### Steps to download and format all the peaks data from ENCODE and Roadmap Epigenomics

1. Download all chromatin feature profiles used for training DeepSEA, specified in Supplementary Table 1 of the DeepSEA manuscript (https://media.nature.com/original/nature-assets/nmeth/journal/v12/n10/extref/nmeth.3547-S2.xlsx). **We have done this for you** (https://zenodo.org/record/2214970/files/chromatin_profiles.tar.gz).
2. For each file, keep the chromosome, start, and end columns. In addition, create a fourth column with the feature’s name. Concatenate all these files and create the distinct features file. **We provide a Python script for this step** (https://github.com/FunctionLab/selene/blob/master/manuscript/case2/1_train_with_online_sampler/data/process_chromatin_profiles.py).
3. Format the data according to the instructions in the “Getting started” tutorial:

a. Sort the file by [chr, start, end]:

~~~
sort -k1V -k2n -k3n **<peak-coordinates-file>** > **<sorted-coordinates-file>**
~~~
b. Compress the file:

~~~
bgzip **<sorted-coordinates-file>**
~~~ This compresses the file to a .gz file in-place. To separately generate the .gz file, run

~~~
bgzip -c **<sorted-coordinates-file> > <sorted-coordinates-file>**.gz
~~~
c. Tabix index the file:

~~~
tabix -p bed **<sorted-coordinates-file>**.gz
~~~ https://github.com/FunctionLab/selene/blob/master/manuscript/case2/1_train_with_online_sampler/data/process_data.sh
4. Download the hg19 FASTA file (https://www.encodeproject.org/files/male.hg19/@@download/male.hg19.fasta.gz).
5. Specify the model architecture, loss, and optimizer as a Python file: https://github.com/FunctionLab/selene/blob/master/selene_sdk/utils/example_model.py
6. Fill out the configuration file with the appropriate file paths and training parameters. We set the training parameters (number of steps, batches, etc.) so that they matched how DeepSEA was originally trained.
7. Run Selene.

#### Case 3: Applying a new model to variants

1. Download the SNPs from the International Genomics of Alzheimer’s Project. (https://www.niagads.org/igap-age-onset-survival-analyses-p-value-only)
2. Group the variants into those with p-values below 0.05 (significant) and those with p-values above 0.50 (nonsignificant).
3. Fill out the configuration file with the paths to the two variants files and the trained model weights file from Case 2.
4. Run Selene.
5. Follow the script provided for this case to analyze the variant predictions (https://github.com/FunctionLab/selene/blob/master/manuscript/case3/2_variant_groups_comparison.sh).

### Code and Data Availability

#### Code

Project homepage: https://selene.flatironinstitute.org

GitHub: https://github.com/FunctionLab/selene

Archived version: https://github.com/FunctionLab/selene/archive/0.2.0.tar.gz

#### Data sources

<u>Cistrome^14^</u>

Cistrome file ID: 33545, measurements from GSM970258 (Xu et al., 2012)

http://dc2.cistrome.org/api/downloads/eyJpZCI6IjMzNTQ1In0%3A1fujCu%3ArNvWLCNoET6o9SdkL8fEv13uRu4b/

<u>ENCODE^19^ and Roadmap Epigenomics^20^ chromatin profiles</u>

Files listed in https://media.nature.com/original/nature-assets/nmeth/journal/v12/n10/extref/nmeth.3547-S2.xlsx

<u>IGAP age at onset survival^15,16^</u>

https://www.niagads.org/datasets/ng00058 (p-values only file)

Processed datasets from these sources are available at the following Zenodo links:

Cistrome:
https://zenodo.org/record/2214130/files/data.tar.gz
ENCODE and Roadmap Epigenomics chromatin profiles:
https://zenodo.org/record/2214970/files/chromatin_profiles.tar.gz
IGAP age at onset survival:
https://zenodo.org/record/1445556/files/variant_effect_prediction_data.tar.gz

## Acknowledgements

The authors acknowledge all members of the Troyanskaya lab for helpful discussions. In addition, the authors thank Dylan Simon for setting up the website and automating updates to the site. The authors are pleased to acknowledge that this work was performed using the high-performance computing resources at Simons Foundation and the TIGRESS computer center at Princeton University. This work was supported by NIH grants R01HG005998, U54HL117798, R01GM071966, and T32HG003284, HHS grant HHSN272201000054C, and Simons Foundation grant 395506. O.G.T. is a CIFAR fellow.

## Contributions

K.M.C and J.Z. conceived the Selene library, K.M.C. and E.M.C. designed, implemented, and documented Selene, K.M.C. performed the analyses described in the manuscript, K.M.C., E.M.C., and O.G.T wrote the manuscript.

## Competing Interests

The authors declare that no competing interests, financial or otherwise, exist.

## References

1. LeCun, Y., Bengio, Y. & Hinton, G. Deep learning. Nature 521, 436–444 (2015).

2. Ching, T., et al. Opportunities and obstacles for deep learning in biology and medicine. J. R. Soc. Interface 15, 20170387 (2018).

3. Segler, M. H. S., Preuss, M. & Waller, M. P. Planning chemical syntheses with deep neural networks and symbolic AI. Nature 555, 604–610 (2018).

4. Zhou, J. & Troyanskaya, O. G. Predicting effects of noncoding variants with deep learning-based sequence model. Nat. Methods 12, 931–934 (2015).

5. Alipanahi, B., Delong, A., Weirauch, M. T. & Frey, B. J. Predicting the sequence specificities of DNA- and RNA-binding proteins by deep learning. Nat. Biotechnol. 33, 831–838 (2015).

6. Kelley, D. R., Snoek, J. & Rinn, J. Basset: Learning the regulatory code of the accessible genome with deep convolutional neural networks. Genome Res. (2016). doi:10.1101/gr.200535.115

7. Angermueller, C., Lee, H. J., Reik, W. & Stegle, O. DeepCpG: accurate prediction of single-cell DNA methylation states using deep learning. Genome Biol. 18, 67 (2017).

8. Kelley, D. R., Reshef, Y., Bileschi, M., Belanger, D., McLean, C. Y. & Snoek, J. Sequential regulatory activity prediction across chromosomes with convolutional neural networks. Genome Res. (2018). doi:10.1101/gr.227819.117

9. Quang, D. & Xie, X. DanQ: a hybrid convolutional and recurrent deep neural network for quantifying the function of DNA sequences. Nucleic Acids Res. 44, e107 (2016).

10. Sundaram, L., et al. Predicting the clinical impact of human mutation with deep neural networks. Nat. Genet. 50, 1161–1170 (2018).

11. Min, S., Lee, B. & Yoon, S. Deep learning in bioinformatics. Brief. Bioinform. 18, 851–869 (2017).

12. Budach, S. & Marsico, A. pysster: Classification Of Biological Sequences By Learning Sequence And Structure Motifs With Convolutional Neural Networks. Bioinformatics (2018). doi:10.1093/bioinformatics/bty222

13. Avsec, Z., et al. Kipoi: accelerating the community exchange and reuse of predictive models for genomics. Preprint at http://dx.doi.org/10.1101/375345 (2018).

14. Mei, S., et al. Cistrome Data Browser: a data portal for ChIP-Seq and chromatin accessibility data in human and mouse. Nucleic Acids Res. 45, D658–D662 (2017).

15. Ruiz, A., et al. Follow-up of loci from the International Genomics of Alzheimer’s Disease Project identifies TRIP4 as a novel susceptibility gene. Transl. Psychiatry 4, e358 (2014).

16. Huang, K.-L., et al. A common haplotype lowers PU.1 expression in myeloid cells and delays onset of Alzheimer’s disease. Nat. Neurosci. 20, 1052–1061 (2017).

17. Li, H., Handsaker, B., Wysoker, A., Fennell, T., Ruan, J., Homer, N., Marth, G., Abecasis, G., Durbin, R. & 1000 Genome Project Data Processing Subgroup. The Sequence Alignment/Map format and SAMtools. Bioinformatics 25, 2078–2079 (2009).

18. Li, H. Tabix: fast retrieval of sequence features from generic TAB-delimited files. Bioinformatics 27, 718–719 (2011).

19. ENCODE Project Consortium. An integrated encyclopedia of DNA elements in the human genome. Nature 489, 57–74 (2012).

20. Roadmap Epigenomics Consortium, Kundaje, A., et al. Integrative analysis of 111 reference human epigenomes. Nature 518, 317–330 (2015).

